# Lung microbiomes’ variable responses to dust exposure

**DOI:** 10.1101/2025.04.10.648223

**Authors:** Mia R. Maltz, Talyssa M. Topacio, David D. Lo, Marina Zaza, Linton Freund, Jon Botthoff, Mark Swenson, David Cocker, Trevor Biddle, Keziyah Yisrael, Diana del Castillo, Ryan Drover, Emma Aronson

**Author notes:** **Correspondence:** Dr. Mia R. Maltz, 1376 Storrs Rd. Unit 4067, Department of Plant Science and Landscape Architecture University of Connecticut, Storrs, CT, 06269-4067 USA. These indicate co-first authors that contributed equally towards experimental manipulations and manuscript preparation (Mia R. Maltz, Talyssa M. Topacio).

## Abstract

Inhalation of dust is significant and relevant to health effects. As pollution and climate change worsen in dryland regions, wind currents entrain loose sediment and dust. This potentially disperses toxic geochemical and microbial burdens throughout the region. When inhaled environmental dust and host-associated microbiomes mingle, they pose exposure risks to host respiratory health. The Salton Sea, California’s largest lake, is shrinking thus exposing nearby communities to playa dust. Therefore, we analyze the effect of Salton Sea dust exposure in murine models to relate lung microbial communities and respiratory health. We used an environmental chamber to expose mice to dust filtrate or ambient air and examined the effects of those exposures on lung microbiomes. We found that lung microbial composition varied by dust exposure. Furthermore, dust elicited neutrophil recruitment and immune responses more than mice exposed to ambient air. Sources of dust differentially affected the composition of the lung core microbiome. Lung microbial diversity correlated with neutrophil recruitment as lungs associated with inflammatory responses harbored more diverse microbiomes. Although Salton Sea dust influences dust microbiomes and prevalent taxa, these responses were variable. The composition of lungs exposed to dust collected further from the Sea was more similar to lungs from ambient air exposures; in contrast, dust collected near the Sea yielded lung microbiomes that clustered further from lungs exposed to ambient air. As lakes continue to dry out, we expect greater public health risks in proximal dryland regions, which may correlate with dust microbial dispersal-related changes to lung microbiomes.

**Importance:** Dust inhalation can lead to health effects, especially when toxic chemicals and microbes mix in with the dust particles. As California’s Salton Sea dries up, it exposes lake bottom sediments to wind, which disperses the dried sediments. To mimic the effect of inhaling Salton Sea dust, we collected and filtered airborne dust to use in exposure experiments with mice in environmental chambers. We predicted that inhaling small dust particles, chemicals, and microbial residues found in this dust would affect mouse respiratory health, or change the microbes found inside their lungs. We found that inhaling dust led to lung inflammation and the dust source influenced the type of microbes found inside mouse lungs. As lakes continue to dry out, we expect greater health risks and changes to lung microbiomes.

**Graphical Abstract:** 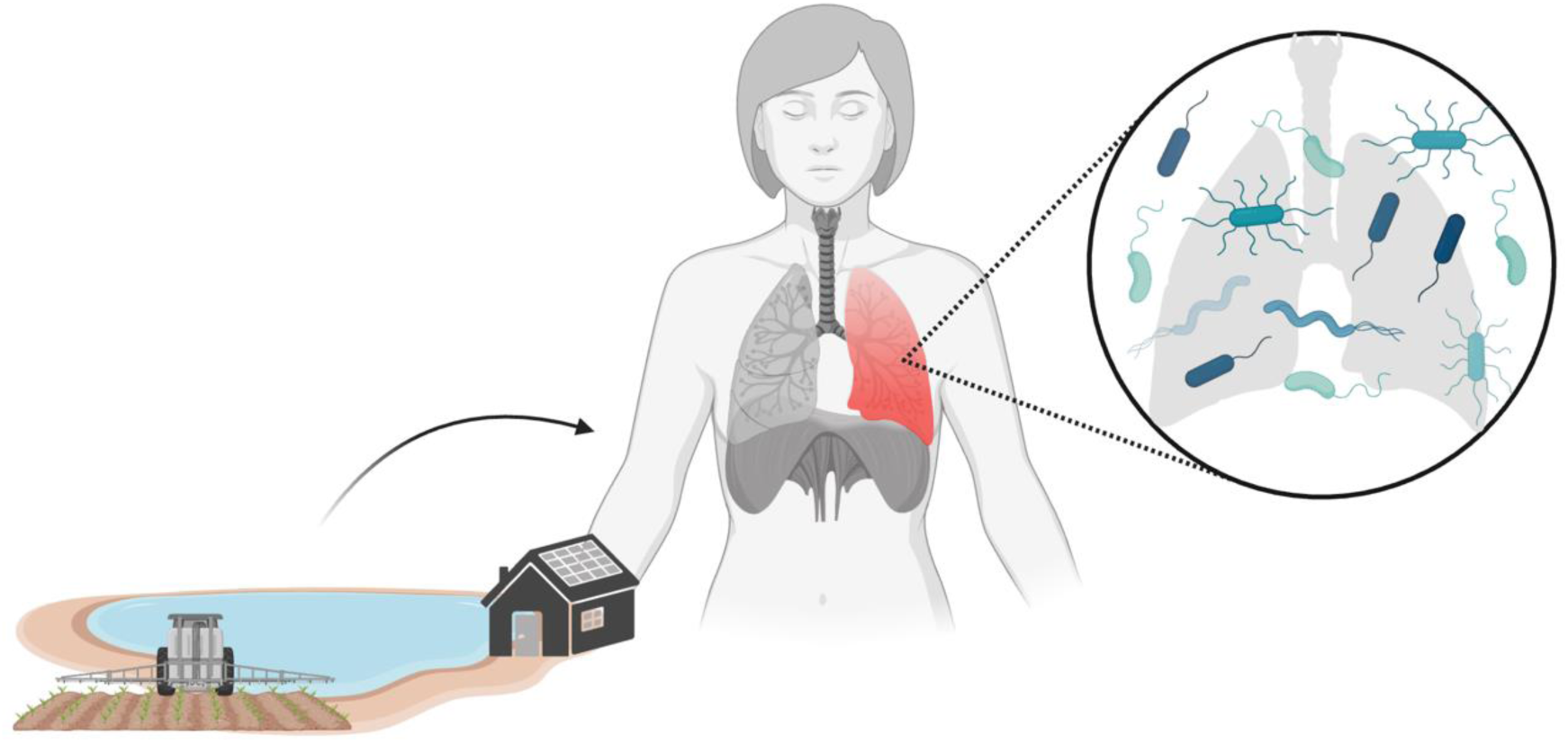

## Introduction

Major dust storms originating in natural desert regions like the Sahara, Australian, and Gobi deserts can carry particulate matter across oceans and continents (1,2). Local dust events that occur in dryland environments and near human settlements are of mounting concern for humans and wildlife. The aeolian environment harbors a variety of airborne particulates composed of both geological and biochemical components, reflective of their provenance (3). At saline terminal lakes, water resource diversion and rising temperatures that exacerbate surface evaporation often lead to waterline recession; this recession may lead to greater sediment (i.e., playa) exposure and increased dust storm frequency (4). Waterline recession and consequential dust events emit insoluble minerals, organic compounds, microbial cells and spores, and microbial exudates, which may become concentrated prior to being aerosolized into local environments. This phenomenon can be harmful to public health; inhaling these increasingly concentrated aerosolized substances presents risks for residents and local communities in surrounding regions (3,5).

At the Salton Sea, a terminal lake located in California’s Imperial Valley, ecological crises have led to a public health concern. The lake was once a popular destination for watersports and tourism, but now the degraded lake basin is notorious for its massive ecological die-offs and toxic marine environment (6). Agricultural waste dumping and a series of freshwater diversions (7) have left the Salton Sea heavily concentrated with unsafe levels of heavy metals, organic residues, and pesticides (8). Increasing regional temperatures and continually decreasing water inputs have led to the lake’s steady recession, transforming the previously submerged lake bottom into newly exposed playa (9). This concentrated, crust-like playa is vulnerable to aerosolization during dust events. Because of these aerosolized exposure risks, the Salton Sea has become a primary suspect for the region’s disproportionately higher rates of asthma-related emergencies, as compared to the total population of California (10,11).

Inhaling dust-derived substances may damage lung health. During normal aspiration, foreign materials from the aeolian environment are consistently introduced into the mammalian respiratory tract (12). Human-host related mechanisms dispel these substances to prevent deep penetration into pulmonary systems. Mucociliary action, antimicrobial peptides, and innate or adaptive immunity contribute to proper clearance of these substances and other potentially harmful agents. However, increased agent frequency or load can overwhelm or bypass these mechanisms, which may result in pulmonary dysbiosis and cause respiratory distress upon inhalation.

In immunocompetent individuals, respiratory tracts are home to low-biomass microbial communities. Despite the tract having once been considered sterile, it is now suspected that microorganisms in the human respiratory tract result from incidental inhalation and micro-aspiration (13). While microbes in the upper respiratory tract (URT) have been found to support host clearance mechanisms and assist in defending against the burden of harmful foreign agents, the dispersion of microbes across the human airway may encourage niche partitioning, owing to either taxonomic, functional, or commensal activities (14). Metabolic and ecophysiological differences among microbial groups, including pathogenicity, as well as the importance of microbiomes for human physiology and disease, underscore the need to consider microbiome composition within the respiratory tract in this system (1,15). Pathogenic potential, timing of microbial arrival, and competition may influence the lung microbiome structure and composition, especially in the lower respiratory tract (LRT), where the steady-state, low-level microbiome must withstand low nutrient supply, stringently high oxygen levels, and antibacterial host responses attributed to the innate immune system.

As the lung microbiome structure correlates with the development and regulation of host immune systems (13), lung microbiomes may directly influence or respond to host pulmonary diseases. Under various disease states, lung microbial composition and diversity shift in patients suffering from asthma, chronic obstructive pulmonary disease (COPD), and cystic fibrosis (CF; (16)). However, it is not clear whether a host’s lung microbiome profile can be used to explicitly differentiate between a healthy and diseased lung phenotype. While pulmonary inflammatory responses may correlate with compositional shifts in lung microbial communities, these responses may be contingent on temporal, climatic, and other external variables, including consistent exposure to environmental aerosols, as suspected among residents in and around the Salton Sea basin.

This study characterizes ambient lung microbiomes as compared to experimentally induced aerosolized dust exposures. We chronically exposed mice to dust collected near California’s Salton Sea, which has been shown to trigger an acute innate immune response within the murine lung (17). Since short exposures to comparable desert dust collected further away from the Sea yielded attenuated immune responses (18), we experimented with dust collected at sites near and further from the Sea to compare exposure impacts on lung microbiomes. Using our novel array of exposure chambers (19), we examined microbial communities in the lungs of mice exposed to dust as compared to mice inhaling filtered air.

Mouse models have been used to understand respiratory health issues because murine lungs can replicate important features of human lung disease pathophysiology; therefore, murine models can be used to observe effects on human lung health, dysbiosis, and responses to medical treatments of pulmonary diseases (20). Focusing on understanding host physiology associated with environmental exposures, prior murine work with C57BL/6 mouse models in this system characterized neurological and immune responses to inhaled vapors and aerosolized substances originating from the Salton Sea region (18,21). While neurological responses to inflammation may vary by mouse sex (22), pulmonary inflammatory responses did not vary by sex. Additionally, lung microbiomes of healthy, C57BL/6 mice may be susceptible to environmentally dependent convergence after at least seven days of co-housing (23). If our chamber experiments with C57BL/6 mice facilitate lung microbiome clustering among animals exposed to filtered air, then we would expect to observe similar compositional clustering among the lung microbiomes of dust-exposed mice. Likewise, if we observe convergent lung microbiome communities, then we hypothesize that there will be no detectable differences observed between right and left or partitioned lobes from the same animals. Moreover, dust exposure would affect mouse health and the composition of their lung microbiomes.

If dust exposure alters health or microbiome status, then baseline lung microbiomes will be affected by chronic exposure to environmental aerosols; furthermore, we hypothesize that these exposures will change lung microbial composition and diversity.

Previous host immunology studies revealed that chronic exposure to dust collected at the Salton Sea at different time points consistently elicits host pulmonary inflammation and other host responses (18), and these inflammatory responses were dissimilar to traditional allergic asthmatic cases. Due to the biochemical changes that occur during lung inflammation, we hypothesize that exposure-induced pulmonary inflammation, regardless of dust variability, will consistently alter lung microbial composition and diversity.

The public health crisis at the Sea corresponds with landscape level changes and increased dust promotion from this degraded ecosystem at the brink of collapse (24–26). The drying lake has the potential to emit toxic substances that may influence public health and host microbiome status. Moreover, as pollution and climate change weaken the resilience of terminal lakes such as the Great Salt Lake, the Aral Sea, and the Salton Sea, it is imperative to understand risks and support improved public health outcomes for inhabitants of these regions.

## Materials and Methods

### Salton Sea dust collection and processing

Passive dust collectors (27) were deployed at two sites of varying distance to the Salton Sea perimeter (Fig. 1). The Palm Desert (PD) site (33°46’25.7“N 116°21’10.3“W) is located 25.5 miles from the lakebed’s nearest waterline and serves as a geographic control site. Wister (WI) is a site (33°17’01.9“N 115°36’00.3“W) situated less than 2 miles off the southeastern edge of the Salton Sea, and dust from this site has consistently yielded histological inflammatory responses and exacerbated pulmonary health statuses in murine systems, as per Biddle et al. (18). We selected two deployment time points: August-October 2020 (WI2020) and September-December 2021 (WI2021, PD2021) because of previous illustrative and immunological analyses. The illustrative analyses showed higher levels of organic matter in Wister dust from these time points, supported by the immunological analyses.

**Figure 1.**
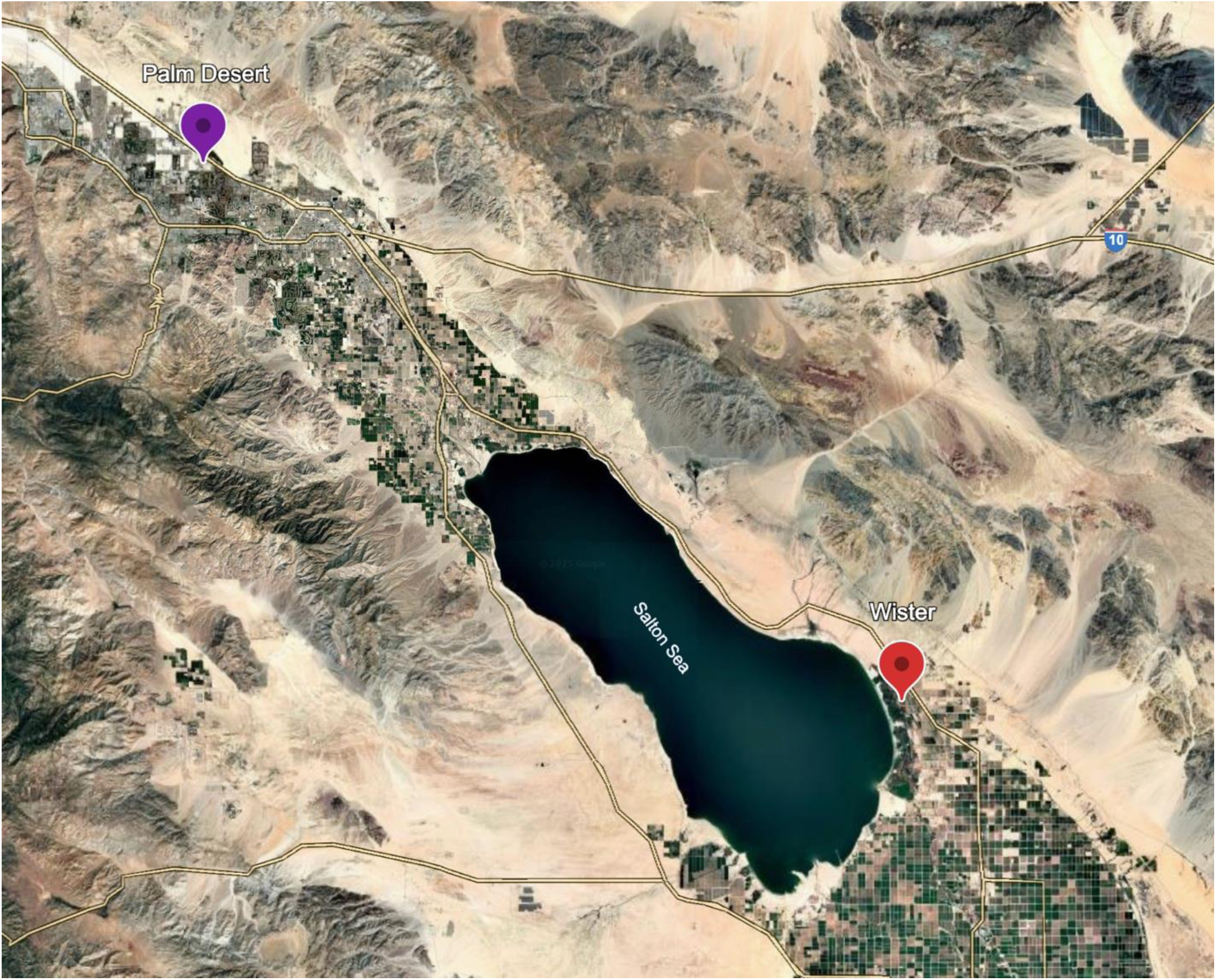
Map of Salton Sea and dust collection sites. Wister (WI, red) is located < 2mi from the eastern edge of the Salton Sea. Palm Desert (PD) is located approximately 26mi northwest from the Salton Sea. Dust material from Boyd Deep Canyon (BDC) was used in previous host pulmonary inflammation studies (18), and is a site located approximately 20mi northwest from the Salton Sea.

After the deployment period, dust collectors were rinsed in MilliQ water and aqueous suspensions were filtered through 0.2 μM mesh. Because the gauge of the mesh was big enough to let water and minerals pass through, but too small to allow for microbial cells to pass through, the remaining lyophilized and concentrated filtrate should be mostly devoid of viable, intact, microbial cells. Subsequently, this concentrated filtrate was resuspended in MilliQ water before aerosolization.

### Animal use ethics

Mice studies were performed in compliance with the University of California, Riverside’s Institutional IACUC and NIH guidelines. Male and female, 8- to 9-week-old C57BL/6 mice were purchased from Jackson Labs (Sacramento, CA). Mice were acclimated for approximately one week in a specific-pathogen free vivarium (University of California, Riverside) before use in our chamber exposure studies.

### Chamber exposures

Cages of 3-4 C57BL/6 mice were randomized between two 540L treatment chambers (exposed and control) modeled after those described in Peng et al. (19). Equal counts of male and female mice were distributed between the two treatment groups. Mice were kept in their respective chambers for 7 days with access to food and water, as needed.

For each experiment, the exposure chamber was filled with dry, filtered air; for the experimental dust manipulations, we mixed this filtered air with aerosolized filtrate suspension passed through a silica drying column as described in Biddle et al. (17). Filtrate concentration was maintained at approximately 1500 μg/m^3^ over the experiment’s entire duration. In contrast, contemporaneous control chambers received dry, filtered air only. These control-air mice were used to define a “baseline” phenotype within the context of this study based on the expectation that this group would exhibit ambient inflammation in comparison to their relative dust-exposed treatment groups at the end of the 7-day exposure period.

### Lower respiratory tract dissection and processing

At the end of the 7-day chamber exposure period, mice were euthanized with isoflurane and cervical dislocation, as per humane animal use protocols. Extraction of LRT tissue was conducted at the mid-trachea for each animal, and lung lobes were separated at the tracheal bifurcation before storing at -80C for downstream use.

Additional lung tissues and bronchoalveolar lavage fluid (BALF) were collected for subsequent analysis of host immune response as described in Biddle et al., 2023 (18). BALF samples were stained with fluorescent antibodies: anti-CD45 FITC (BioLegend, San Diego, USA; Clone 30-F11), anti-CD19 PE-Cy5 (eBioscience, San Diego, USA; Clone MB19-1), anti-CD3 Alexa Fluor 700 (BioLegend, San Diego, USA; Clone 17A2), anti-Ly6G BV510 (BioLegend, San Diego, USA; Clone 1A8), anti-CD11b BV421 (BioLegend, San Diego, USA; Clone M1/70), anti-CD11c PE-Cy7 (BioLegend, San Diego, USA; Clone N418) and anti-SiglecF APC (BioLegend, San Diego, USA; Clone S17007L). Flow cytometry was performed on a MoFlo Astrios (Beckman Coulter, Carlsbad, CA) and gating and analysis were done using FlowJo (Version 10.71).

### Lung microbiome library prep and sequencing

Microbial DNA from whole- or half-lung lobes was extracted using a HostZERO Microbial DNA extraction kit (Zymo Research, Irvine, CA) following a modified version of the manufacturer’s protocol for solid tissue samples by disrupting these microbe-containing lung tissues. We performed tissue lysis using 2.0 mm beads on a MP Bio Fastprep Classic (MP Biomedicals, Irvine, CA) for one minute at maximum speed, followed by a 2-minute centrifugation cycle to separate eukaryotic and microbial components. After two subsequent cycles of mixing and centrifugation, we resuspended pellets and performed three incubations; the first was at 37 °C for 30 minutes, the second included proteinase K treatments at 55 °C for 10 minutes, and the third incubation was at room temperature for 5 minutes. Next, we transferred solutions to lysis tubes containing 0.1 and 0.5 mm beads for five cycles of lysis and 5 minute incubations on ice, followed by centrifugation and downstream DNA extraction, as per the manufacturer’s recommendation. However, we modified this protocol to maximize the microbial extraction process by transferring a total of 700 μl of supernatant into two parallel extractions from each of our whole- or half-lung samples, which we subsequently combined as a singular DNA template corresponding to that particular animal.

Due to the lungs being characterized by low microbial biomass, negative controls were used alongside DNA extractions to control for potential contaminants in downstream analyses. DNA extracts were quantified with Qubit (Invitrogen, Carlsbad, CA); double-stranded DNA mass (ng) concentrations from negative control samples were undetectable (per μl) in comparison to template from lung samples (23). Extracts with detectable DNA concentrations were sent to Zymo Research (Irvine, CA) for targeted-amplicon library preparation of 16S V3-V4 rRNA gene sequencing.

Prior to library preparation, a High Resolution Cleanup (HRC) PCR inhibitor removal step was conducted using the OneStep PCR Inhibitor Removal Kit (Zymo Research, Irvine, CA). Following this step, 16S rRNA amplification was done for the V3 and V4 regions using the Quick-16S NGS Library Prep Kit (Zymo Research, Irvine, CA) with added Peptide Nucleic Acids (PNAs; i.e., mitochondrial blockers), as per Lundberg et al. (28), to minimize the amplification of eukaryotic mitochondrial DNA. Negative controls were maintained throughout the amplification pipeline for tracking contamination and protocol efficacy. Libraries were sequenced with an Illumina NextSeq 2000 using the p1 reagent kit (600 cycles) and a 30% PhiX spike to promote read diversity. Sequences were submitted to the National Center for Biotechnology Information Sequence Read Archive under BioProject PRJNA1124545.

### Bioinformatics amplicon sequence analysis

16S rRNA (V3 and V4) amplicon sequence data were analyzed using methods described in Freund, et al. (29,30). Sequence quality was assessed using FastQC and eestats2 (31). Sequences that passed through these quality thresholds were trimmed and ASVs were assigned using the DADA2 pipeline (32). The R “decontam” package was used to identify and remove amplicon sequencing variants (ASVs) associated with negative control samples and potential contaminants, as well as chloroplast or mitochondria-associated taxa.

### Data analysis and statistics

All 16S rRNA amplicon sequencing data were analyzed in RStudio (R software version 3.18) using methods described in Freund et al. (2025). Normal distribution for Shannon-Weiner diversity and species richness among the lung microbiomes of exposure treatment groups were determined using the Shapiro-Wilks test. Since Shannon-Weiner diversity (*P*<.001) and species richness (*P*<.001) were not normally distributed, we used the Kruskal-Wallis test (“kruskal.test” function, “stats” package) to compare variance of means for both Shannon-Weiner diversity and species richness between exposure treatment groups. If the variance within the means of Shannon-Weiner diversity or species richness differed significantly between groups (α=0.05), we performed a Dunn test (“dunn_test” function, “rstatix” package) to determine which pair(s) of exposure treatment groups differed in Shannon-Weiner diversity or taxa richness.

The R “vegan” package was used for beta diversity. Data was transformed by center-log ratio (CLR) with the “decostand” function, and a Principal Coordinates Analysis (PCoA) was performed and visualized on Aitchison distances (33–35). Before pairwise comparisons were run, “betadisper” was used to assess homogeneity of variance. After ensuring that dispersion did not differ significantly between groups, permutational multivariate analysis of variance (PERMANOVA) was performed using “adonis2,” and comparisons between single pairings were clarified with “pairwiseAdonis” (36).

## Results

### Host inflammatory response, lung microbiome composition, and dust exposure

We found that exposure to environmental dust elicits significant neutrophilic inflammation in mouse lungs when compared to their contemporaneous, ambient air control group (Fig. 2). Using a Kruskal-Wallis test for non-normally distributed data, we found that average neutrophil recruitment as a percentage of CD45+ cells differed significantly among treatment groups (P=0.004). A Wilcoxon test verified significant increases in neutrophil percentages in PD2021 dust (P=0.029) and WI2020 dust (P=0.029) exposed mice, and a marginally significant increase in neutrophil recruitment in animals exposed to WI2021 dust (P=0.057) as compared to their contemporaneous controls. For mice exposed to WI2020 dust, average neutrophil recruitment was significantly higher than in mice exposed to PD2021 dust (P=0.029) or WI2021 dust (P=0.029), indicating a higher magnitude of neutrophilic pulmonary inflammation elicited by dust collected in fall of 2020 at the Wister site (close to the Salton Sea; WI2020). In contrast, no significant differences in eosinophil recruitment were observed among dust-exposure groups and their contemporaneous controls.

**Figure 2.**
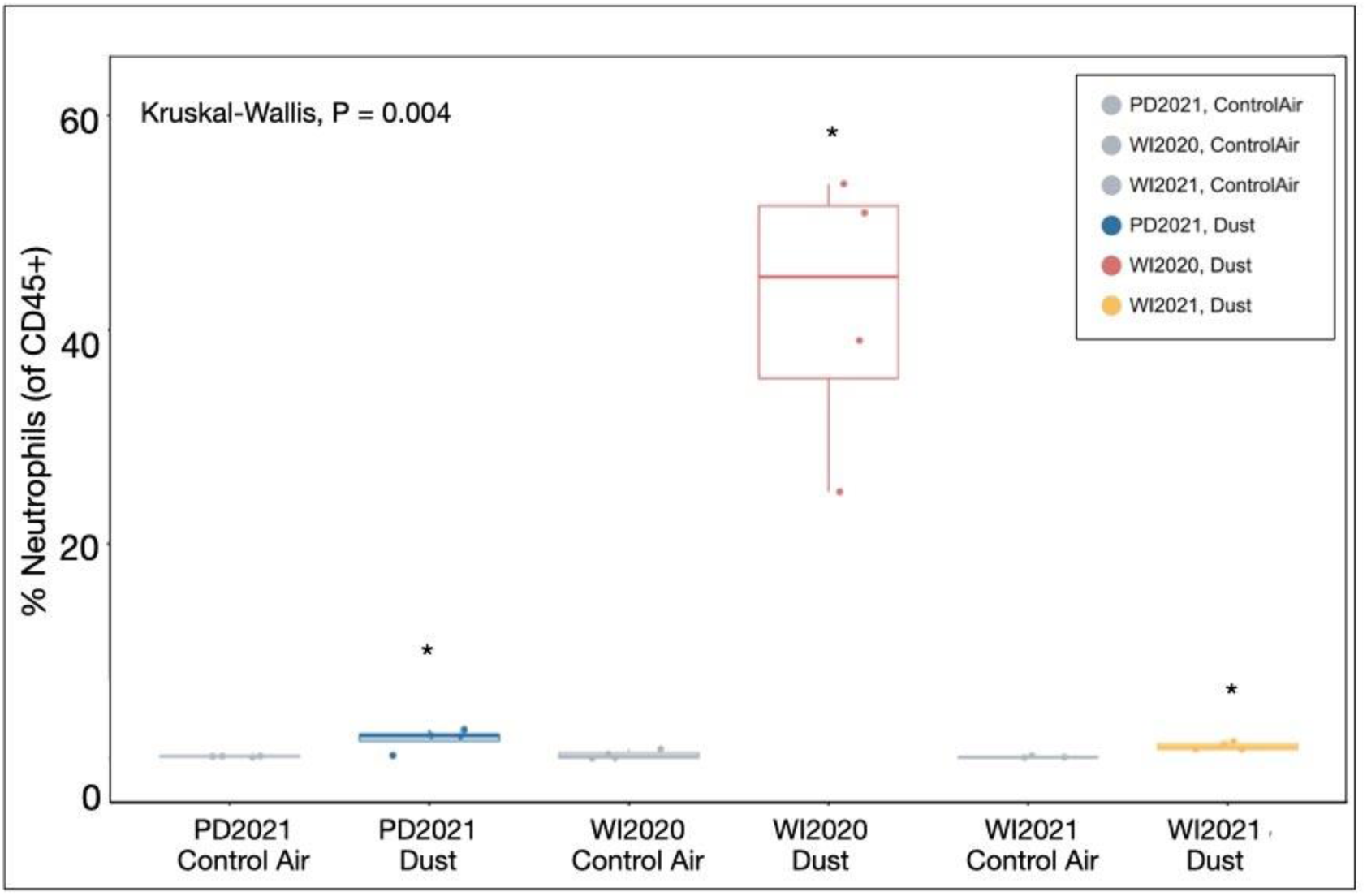
Inflammatory responses to dust exposure. Average neutrophil recruitment (% of CD45+ cells) in mouse lungs differed significantly by exposure material (P=0.004). Neutrophil recruitment was significantly higher in dust-exposed lungs when compared to their contemporaneous controls.

In terms of host neutrophil recruitment, control-air-exposed mice were deemed healthy, and were used to characterize a baseline lung microbiome within a shared environment. Using Aitchison distance matrices, we used a Principal Coordinates Analysis (PCoA) to visualize dissimilarities in lung microbial community composition by treatment group, such as exposure treatment (Fig. 3). Lung microbiomes of PD2021-, WI2020-, and WI2021-exposed mice have more dispersion than control-air-exposed mice. Moreover, the lung microbiomes of mice exposed to PD2021 dust cluster closely to mice exposed to ambient air, while microbiomes of mice exposed to WI2020 and WI2021 dust cluster further away from this group and each other. A PERMANOVA revealed that lung microbiome community composition significantly differed between exposure treatment groups (P<0.001). Specifically, exposure to WI2021 dust significantly changed lung microbiome composition in comparison to their contemporaneous control-air microbiomes (P=0.002). While exposure to PD2021 dust did not significantly change lung microbiome composition in comparison to its contemporaneous control group (P=0.630), the composition of PD-exposed dust lung microbiomes differed significantly from WI2020 dust (P=0.003) and WI2021 dust (P=0.002) microbiomes. Likewise, lung microbiome composition in the WI2020-exposed group differed significantly from lung microbiomes associated with the WI2021-exposed group (P=0.002).

**Figure 3.**
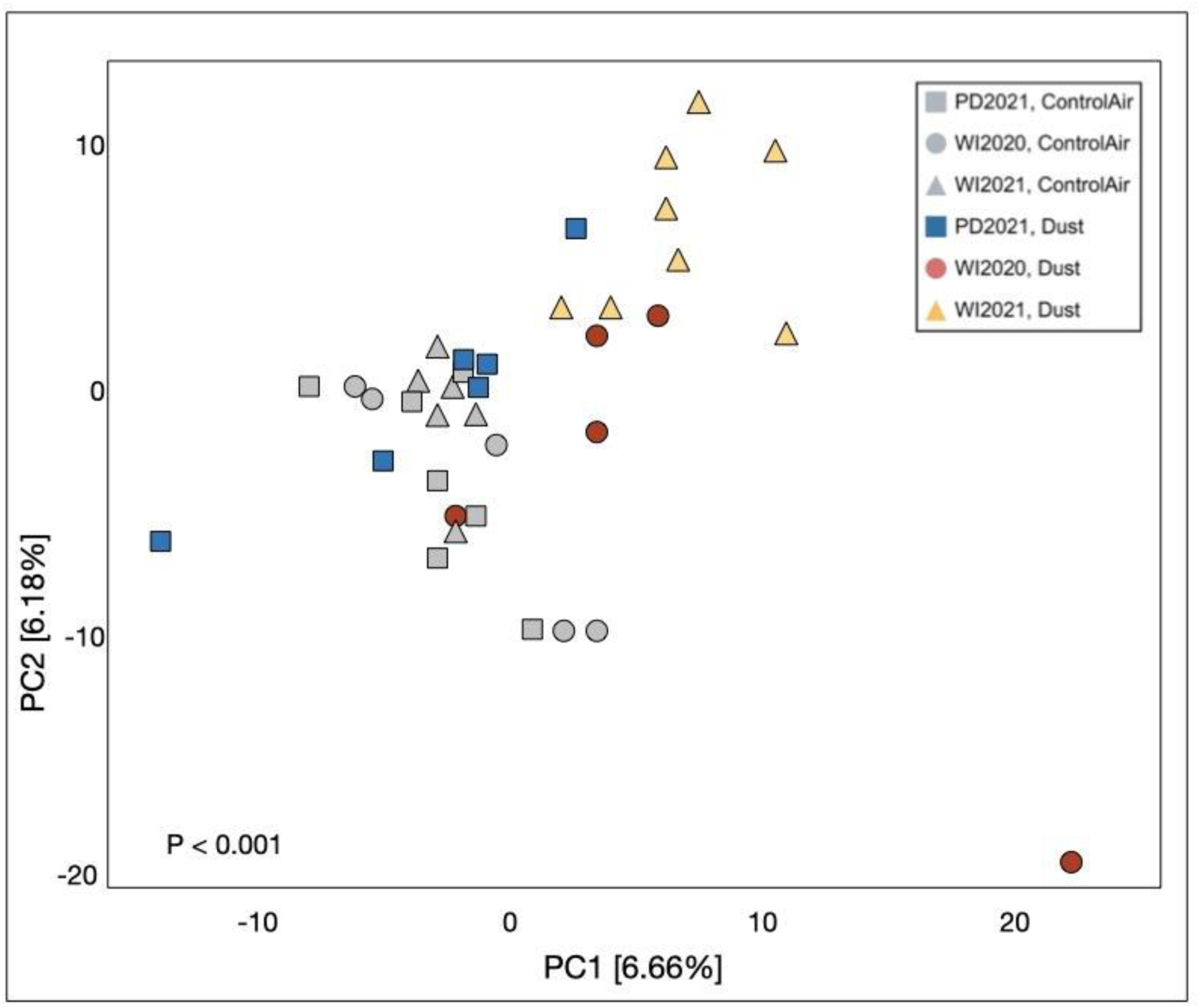
Microbial community composition in exposed mouse lungs. Lung microbial community composition differs significantly among exposure material groups (P<0.001). Microbiomes of mice exposed to PD2021 dust cluster closely to mice exposed to ambient air, while microbiomes of mice exposed to WI2020 and WI2021 dust cluster separately from this group and each other. Overall, dust-exposed microbiomes are more widely dispersed than controls.

### Baseline lung microbial diversity and dust exposure

We calculated species richness and Shannon-Weiner diversity of lung microbial communities and used Kruskal-Wallis tests for determining differences between treatment groups (Fig. 4). Species richness did not differ significantly between groups (P=0.26). Likewise, Shannon-Weiner diversity did not differ significantly in dust-exposed lung microbiomes when compared to their contemporaneous ambient air controls (PD2021 P=0.95, WI2020 P=0.078, WI2020 P=0.23) When comparing Shannon-Weiner diversity between dust-exposed groups, lung microbiome diversity in WI2020-exposed mice did not differ significantly from that of PD2021-exposed mice (P=0.21). In contrast, Shannon-Weiner diversity differed significantly between WI2020- and WI2021-exposed mice (P<0.001), ostensibly because of variability in evenness across these two Wister-dust-exposed groups.

**Figure 4.**
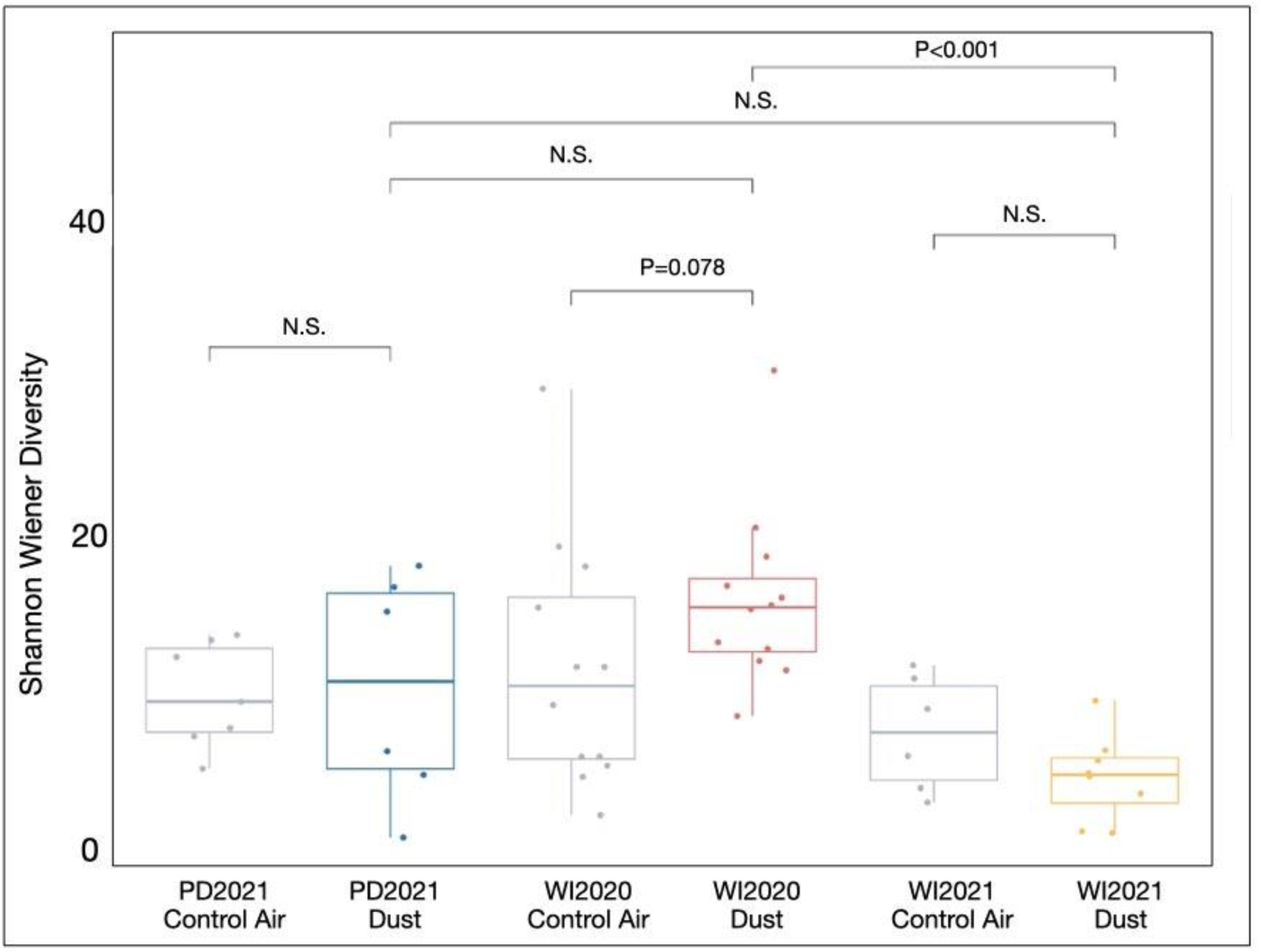
Microbial diversity varies by dust exposure. While average Shannon-Weiner diversity differs significantly among exposure groups (P=0.0027), dust exposures do not differ significantly from their contemporaneous controls. Average Shannon-Weiner diversity is significantly higher in WI2020-exposed lung microbiomes compared to WI2021-exposed lung microbiomes.

We used core microbiome analysis (37) to visualize evenness and taxa prevalence in each treatment group at a minimum threshold of 1% relative abundance (RA; Fig. 5). Species evenness was significantly different among groups according to exposure material (P<0.01) with WI2021-exposed lung microbiomes being significantly less even compared to control air microbiomes (P=0.016) and WI2020-exposed lung microbiomes (P=0.019). When comparing lung microbiome taxa abundance between control-air- and dust-exposed mice, the WI2020 group displayed higher taxa abundance, with more or different taxa being uniquely present or more prevalent at 1% RA. These taxa include *Achromobacter*, *Atopostipes, Enhydrobacter, Methylobacterium, Methylorubrum,* and *Microbacterium*. In contrast, two taxa, *Paenarthrobacter* and *Staphylococcus,* were prevalent among 75-100% of WI2021 lung microbiome samples. Moreover, the WI2021-exposed group displayed lower abundances across core taxa, with most taxa being prevalent at approximately 25% of sampled lungs.

**Figure 5.**
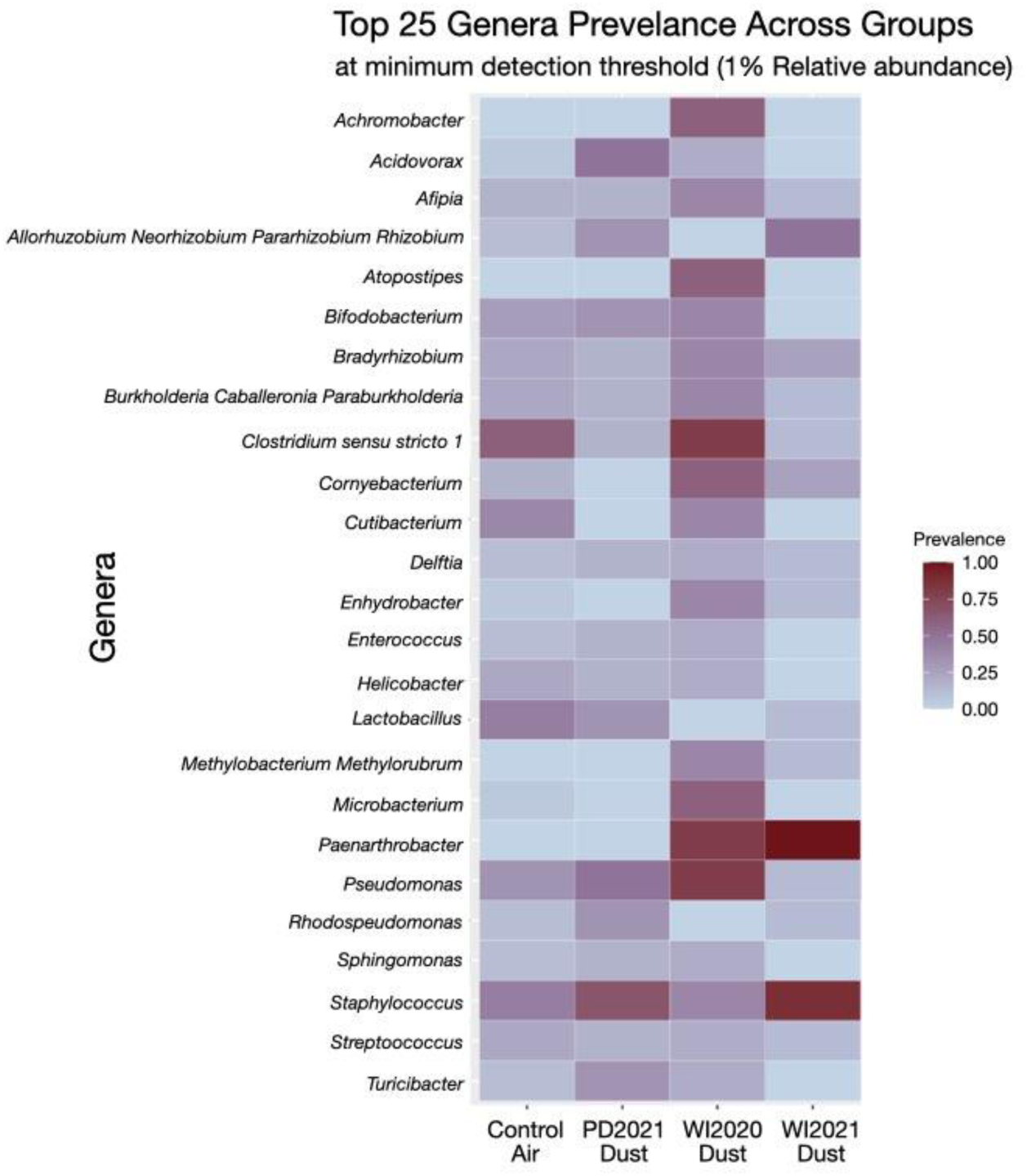
Prevalent microbial genera across treatment groups. Prevalence is calculated as a percentage of the samples where the taxa are observed at 1% relative abundance or higher. In control air-exposed lung microbiomes, most taxa are moderately prevalent (25-75%). In WI2021-exposed lung microbiomes, few taxa are represented at high (>75%) prevalence, while many other taxa are represented at low (<25%) prevalence.

We used PERMANOVA to assess if differences in lung microbiome responses following chronic dust exposure could be attributed to mouse sex or lung lobe section (left vs. right lobe). We did not detect any effects due to sex or lobes. Among control-air-exposed mice only, no significant differences in baseline lung microbiome composition were attributed to either sex (P=0.567) or lobe section (P=0.721). Similarly, among dust-exposed mice only, no significant differences were detected in lung microbiome composition between sexes (P=0.848) or lobe sections (P=0.906).

## Discussion

In this study, we showed that lung microbial community composition varied by dust exposure. Neutrophil recruitment was found to be higher in the lungs of dust-exposed mice compared to mice exposed to ambient filtered air. We characterized a baseline lung microbiome among mice used in this study; however, the specific microbiome here described is not broadly indicative of host pulmonary health or inflammatory state beyond the scope of this study. While no differences in lung microbial communities were detected by mouse sex or lung lobe, the core lung microbiome diversity and composition were affected by dust source and exposure. Indeed, mice exposed to dust collected at the Wister site, which elicited a heightened neutrophilic immune response were characterized by higher beta- and alpha-diversity and evenness in their lung microbiomes than were Wister dust-exposed mice exhibiting lesser neutrophil recruitment. Likewise, dust from Wister exerts an influence on dust microbial composition and changes the relative abundance of prevalent taxa. Mouse lungs exposed to dust from Palm Desert, which is further from the Sea, were more similar to lungs from ambient control air exposures. Conversely, exposures to dust collected near the Sea from the Wister site in 2021 yielded lung microbiomes that clustered further from lungs exposed to ambient air.

Neutrophil recruitment differed between control and exposed groups, which suggests that chronic dust exposure does elicit neutrophilic inflammation; however, the magnitude of that inflammation is not consistent with the changes in lung microbiome diversity and composition. This suggests that the host lung microbiome is responding directly to dust exposure rather than the physiological effects of lung inflammation. It is unclear if changes to lung architecture during disease drive lung microbial dysbiosis, or if dysbiosis instigates inflammation. Previous studies suggest that the lung microbiome profile and its relationship to the host’s immune response is contingent on endotype (38). One study observed a lack of relationship between Type 2 asthma inflammatory markers and lung microbiome composition but went on to suggest that dysbiosis in the lung microbiome could be highly relevant in patients with severe, non-Type 2 asthma (characterized by neutrophil-dominant inflammation), akin to what we have observed in this study (39). Furthermore, another study observed significantly higher bacterial species richness in asthma patient sputum samples characterized by mixed (neutrophil and eosinophil) or neutrophil-dominant endotypes when compared to eosinophil-dominant and paucigranulocytic endotypes (40). In our study, chronic exposure to abiotic dust material led to significant neutrophil-dominant inflammation when compared to their contemporaneous ambient air controls. Upon referencing significant changes observed in lung microbiome composition (Fig. 3), our findings suggest that dust-induced dysbiosis in the lung microbiome resulting from chronic exposure may be driving neutrophil recruitment and thus triggering significant pulmonary inflammation.

Host inflammatory response and lung microbiome composition are independently affected by chronic dust exposure. Previous studies show that low microbial diversity is related to poor lung function (41–44). Likewise, lung microbiomes in children with asthma may be composed differently than healthy children’s lungs (45), which could relate to disease severity or type of medical intervention. Our study found that while chronic dust exposure altered lung microbiome diversity, it did so independently from host pulmonary inflammatory state.

Although microbial taxa richness did not vary, baseline lung microbiome diversity changed from chronic exposure to Wister dust. Mouse lungs exposed to dust from Wister collected in 2020 were as diverse as Palm Desert (2021)-exposed mouse lungs. However, the mouse lungs exposed to dust from Wister in 2021 surpassed microbial diversity levels of the other treatment groups. Given that dust storms entrain microbial residues, chemical constituents, and particulates that could result in host responses to that dust, the presence or prevalence of these substances likely changes the relative abundance of taxa found in lung microbiomes. Previous studies show that exposure to particulate matter, dust, smoke, or ozone can lead to lung microbial dysbiosis, which may be related to asthmatic inflammation or other respiratory pathologies (46,47). Yet data from several of these studies are correlative and are apt to raise more questions about whether the substances alter lung microbiomes directly or lung injury following exposure to these types of residues or substances alters the respiratory ecosystem, thereby impacting the microbiome indirectly. Our study showed higher evenness in mouse lungs exposed to dust from Wister in 2020 than other dust sources or dates, which contributed to the heightened diversity detected in these lung microbiomes.

We detected higher prevalence of Gram-positive bacteria *Paenarthrobacter* and *Staphylococcus* in lungs exposed to dust from Wister collected in 2021, which were common across animals in that treatment group and correlated with lower neutrophil recruitment. In contrast, the lungs exposed to dust from Wister in 2020 had higher neutrophil recruitment and more Gram-negative *Pseudomonas* spp. than lungs exposed to WI2021 dust. In contrast to our finding that lungs enriched in *Pseudomonas* harbored high microbial diversity, previous work on *Pseudomonas aeruginosa* found correlations with *Pseudomonas* and decreased lung microbial diversity (48–50). *Pseudomonas* has been commonly found in patients’ lungs with severe asthmatic inflammation (51,52), yet *Pseudomonas* is virtually undetectable in normal healthy lungs. Other studies suggest that neutrophil recruitment and host inflammatory response may generally relate to Gram-negative bacteria, like *Pseudomonas,* that are enriched with lipopolysaccharides (LPS) and mediate neutrophil activation. Likewise, our findings showed more prevalent Gram-negative taxa in lung microbial communities exposed to WI2020 dust than were found in WI2021-exposed lungs. While different diseases, including asthma, may alter lung microbiome structure and function, other demographic features (such as age and gender) could determine whether exposed residents in the Salton Sea region would be susceptible to cardiopulmonary or neurological disease (53–55). Our study suggests that exposure to dust could influence lung microbiome structure in vulnerable populations.

Overall, our findings show that environmental dust exposure changes lung microbial communities. Given the predicted increase in Salton Sea dust emissions, entrained dust may introduce contaminants, minerals, and other potential toxins into the atmosphere. These changing conditions may also influence the geographic distribution and functional capacity of aeolian microbial community originating from the drying Salton Sea lakebed. However, our study utilized filtered, abiotic dust material and thus, does not represent the direct effects of the Salton Sea aeolian microbiome on host lung microbiome composition. Altogether, microbial residues, chemical constituents, along with mineral and organic particulates in local dust emissions can compound exposure risks for the region’s inhabitants, as well as have implications for ecosystem stability, conservation, and public health.

## Supplementary Figures

**S1.**
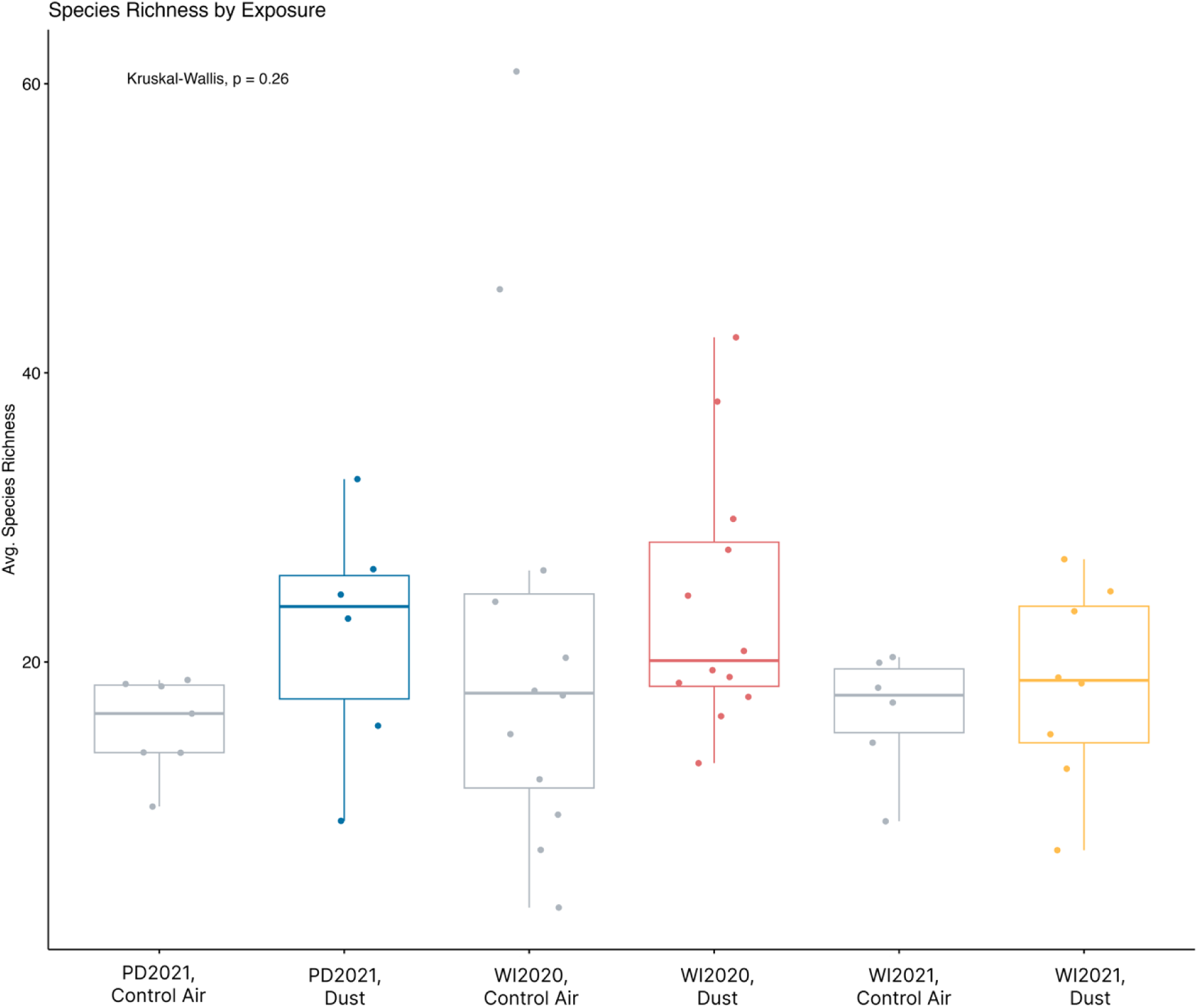
Microbial species richness and dust exposure. Average species richness does not differ significantly among exposure groups (P=0.26).

## References

1. Maltz MR, Carey CJ, Freund HL, Botthoff JK, Hart SC, Stajich JE, et al. Landscape Topography and Regional Drought Alters Dust Microbiomes in the Sierra Nevada of California. Front Microbiol. 2022 Jun 28;13:856454.

2. Yamaguchi N, Park J, Kodama M, Ichijo T, Baba T, Nasu M. Changes in the Airborne Bacterial Community in Outdoor Environments following Asian Dust Events. Microbes Environ. 2014;29(1):82–8.

3. Esmaeil N, Gharagozloo M, Rezaei A, Grunig G. Dust events, pulmonary diseases and immune system. 2014 Mar 15;

4. Zucca C, Middleton N, Kang U, Liniger H. Shrinking water bodies as hotspots of sand and dust storms: The role of land degradation and sustainable soil and water management. CATENA. 2021 Dec;207:105669.

5. Cook AG, Weinstein P, Centeno JA. Health Effects of Natural Dust: Role of Trace Elements and Compounds. Biol Trace Elem Res. 2005;103(1):001–16.

6. Cohen MJ, Morrison JI, Glenn EP. The Ecology and Future of the Salton Sea. 1999 Feb;

7. Johnston JE, Razafy M, Lugo H, Olmedo L, Farzan SF. The disappearing Salton Sea: A critical reflection on the emerging environmental threat of disappearing saline lakes and potential impacts on children’s health. Sci Total Environ. 2019 May;663:804–17.

8. Moreau MF, Surico-Bennett J, Vicario-Fisher M, Gerads R, Gersberg RM, Hurlbert SH. Selenium, arsenic, DDT and other contaminants in four fish species in the Salton Sea, California, their temporal trends, and their potential impact on human consumers and wildlife. Lake Reserv Manag. 2007 Dec;23(5):536–69.

9. Jones BA, Fleck J. Shrinking lakes, air pollution, and human health: Evidence from California’s Salton Sea. Sci Total Environ. 2020 Apr;712:136490.

10. Farzan SF, Razafy M, Eckel SP, Olmedo L, Bejarano E, Johnston JE. Assessment of Respiratory Health Symptoms and Asthma in Children near a Drying Saline Lake. Int J Environ Res Public Health. 2019 Oct 11;16(20):3828.

11. Miao Y, Porter WC, Schwabe K, LeComte-Hinely J. Evaluating health outcome metrics and their connections to air pollution and vulnerability in Southern California’s Coachella Valley. Sci Total Environ. 2022 May;821:153255.

12. Whiteson KL, Bailey B, Bergkessel M, Conrad D, Delhaes L, Felts B, et al. The Upper Respiratory Tract as a Microbial Source for Pulmonary Infections in Cystic Fibrosis. Parallels from Island Biogeography. Am J Respir Crit Care Med. 2014 Jun 1;189(11):1309–15.

13. Marimón JM. The Lung Microbiome in Health and Respiratory Diseases. Clin Pulm Med. 2018 Jul;25(4):131–7.

14. Hamm PS, Taylor JW, Cook JA, Natvig DO. Decades-old studies of fungi associated with mammalian lungs and modern DNA sequencing approaches help define the nature of the lung mycobiome. Sheppard DC, editor. PLOS Pathog. 2020 Jul 30;16(7):e1008684.

15. Blaser MJ. The microbiome revolution. J Clin Invest. 2014 Oct 1;124(10):4162–5.

16. Dickson RP, Erb-Downward JR, Martinez FJ, Huffnagle GB. The Microbiome and the Respiratory Tract. Annu Rev Physiol. 2016 Feb 10;78(1):481–504.

17. Biddle TA, Li Q, Maltz MR, Tandel PN, Chakraborty R, Yisrael K, et al. Salton Sea aerosol exposure in mice induces a pulmonary response distinct from allergic inflammation. Sci Total Environ. 2021 Oct;792:148450.

18. Biddle TA, Yisrael K, Drover R, Li Q, Maltz MR, Topacio TM, et al. Aerosolized aqueous dust extracts collected near a drying lake trigger acute neutrophilic pulmonary inflammation reminiscent of microbial innate immune ligands. Sci Total Environ. 2023 Feb;858:159882.

19. Peng X, Maltz MR, Botthoff JK, Aronson EL, Nordgren TM, Lo DD, et al. Establishment and characterization of a multi-purpose large animal exposure chamber for investigating health effects. Rev Sci Instrum. 2019 Mar 1;90(3):035115.

20. Rydell-Törmänen K, Johnson JR. The Applicability of Mouse Models to the Study of Human Disease. In: Bertoncello I, editor. Mouse Cell Culture [Internet]. New York, NY: Springer New York; 2019 [cited 2025 Mar 5]. p. 3–22. (Methods in Molecular Biology; vol. 1940). Available from: http://link.springer.com/10.1007/978-1-4939-9086-3_1

21. Llamas MJV. Adult mice exposed to aerosolized Alternaria exhibit neuroinflammation in the brainstem but not rest of brain. 2016.

22. Paolicelli RC, Sierra A, Stevens B, Tremblay ME, Aguzzi A, Ajami B, et al. Microglia states and nomenclature: A field at its crossroads. Neuron. 2022 Nov;110(21):3458–83.

23. Dickson RP, Erb-Downward JR, Falkowski NR, Hunter EM, Ashley SL, Huffnagle GB. The Lung Microbiota of Healthy Mice Are Highly Variable, Cluster by Environment, and Reflect Variation in Baseline Lung Innate Immunity. Am J Respir Crit Care Med. 2018 Aug 15;198(4):497–508.

24. Freund H, Maltz MR, Swenson MP, Topacio TM, Montellano VA, Porter W, et al. Microbiome interactions and their ecological implications at the Salton Sea. Calif Agric. 2022 Apr;76(1):16–26.

25. Miao Y, Porter WC, Benmarhnia T, Lowe C, Lyons TW, Hung C, et al. Source-specific acute cardio-respiratory effects of ambient coarse particulate matter exposure in California’s Salton Sea region. Environ Res Health. 2025 Mar 1;3(1):015006.

26. Sinclair RG, Gaio J, Huazano SD, Wiafe SA, Porter WC. A Balloon Mapping Approach to Forecast Increases in PM10 from the Shrinking Shoreline of the Salton Sea. Geographies. 2024 Oct 17;4(4):630–40.

27. Aciego SM, Riebe CS, Hart SC, Blakowski MA, Carey CJ, Aarons SM, et al. Dust outpaces bedrock in nutrient supply to montane forest ecosystems. Nat Commun. 2017 Mar 28;8(1):14800.

28. Lundberg DS, Yourstone S, Mieczkowski P, Jones CD, Dangl JL. Practical innovations for high-throughput amplicon sequencing. Nat Methods. 2013 Oct;10(10):999–1002.

29. Freund L, Hung C, Topacio TM, Diamond C, Fresquez A, Lyons TW, et al. Diversity of sulfur cycling halophiles within the Salton Sea, California’s largest lake. BMC Microbiol. 2025 Mar 6;25(1):120.

30. Maltz MR, Allen MF, Phillips ML, Hernandez RR, Shulman HB, Freund L, et al. Microbial community structure in recovering forests of Mount St. Helens. Front Microbiomes. 2024 Nov 4;3:1399416.

31. Andrews S. Andrews S. FastQC: A Quality Control Tool for High Throughput Sequence Data. 2010. [Internet]. 2010. Available from: https://www.bioinformatics.babraham.ac.uk/projects/fastqc/

32. Callahan BJ, McMurdie PJ, Rosen MJ, Han AW, Johnson AJA, Holmes SP. DADA2: High-resolution sample inference from Illumina amplicon data. Nat Methods. 2016 Jul;13(7):581–3.

33. Lubbe S, Filzmoser P, Templ M. Comparison of zero replacement strategies for compositional data with large numbers of zeros. Chemom Intell Lab Syst. 2021 Mar;210:104248.

34. Quinn TP, Erb I, Richardson MF, Crowley TM. Understanding sequencing data as compositions: an outlook and review. Wren J, editor. Bioinformatics. 2018 Aug 15;34(16):2870–8.

35. Quinn TP, Erb I, Gloor G, Notredame C, Richardson MF, Crowley TM. A field guide for the compositional analysis of any-omics data. GigaScience. 2019 Sep 1;8(9):giz107.

36. Martinez Arbizu P. pairwiseAdonis [Internet]. 2020. Available from: https://github.com/pmartinezarbizu/pairwiseAdonis

37. Lahti L, Shetty S. Core microbiome [Internet]. 2018. Available from: https://microbiome.github.io/tutorials/Core.html

38. Natalini JG, Singh S, Segal LN. The dynamic lung microbiome in health and disease. Nat Rev Microbiol. 2023 Apr;21(4):222–35.

39. Huang YJ, Charlson ES, Collman RG, Colombini-Hatch S, Martinez FD, Senior RM. The Role of the Lung Microbiome in Health and Disease. A National Heart, Lung, and Blood Institute Workshop Report. Am J Respir Crit Care Med. 2013 Jun 15;187(12):1382–7.

40. Son JH, Kim JH, Chang HS, Park JS, Park CS. Relationship of Microbial Profile With Airway Immune Response in Eosinophilic or Neutrophilic Inflammation of Asthmatics. Allergy Asthma Immunol Res. 2020;12(3):412.

41. Coburn B, Wang PW, Diaz Caballero J, Clark ST, Brahma V, Donaldson S, et al. Lung microbiota across age and disease stage in cystic fibrosis. Sci Rep. 2015 May 14;5(1):10241.

42. Cox MJ, Allgaier M, Taylor B, Baek MS, Huang YJ, Daly RA, et al. Airway Microbiota and Pathogen Abundance in Age-Stratified Cystic Fibrosis Patients. Ratner AJ, editor. PLoS ONE. 2010 Jun 23;5(6):e11044.

43. Flight WG, Smith A, Paisey C, Marchesi JR, Bull MJ, Norville PJ, et al. Rapid Detection of Emerging Pathogens and Loss of Microbial Diversity Associated with Severe Lung Disease in Cystic Fibrosis. McAdam AJ, editor. J Clin Microbiol. 2015 Jul;53(7):2022–9.

44. Hahn A, Warnken S, Pérez-Losada M, Freishtat RJ, Crandall KA. Microbial diversity within the airway microbiome in chronic pediatric lung diseases. Infect Genet Evol. 2018 Sep;63:316–25.

45. Hilty M, Burke C, Pedro H, Cardenas P, Bush A, Bossley C, et al. Disordered Microbial Communities in Asthmatic Airways. Neyrolles O, editor. PLoS ONE. 2010 Jan 5;5(1):e8578.

46. Hosang L, Canals RC, Van Der Flier FJ, Hollensteiner J, Daniel R, Flügel A, et al. The lung microbiome regulates brain autoimmunity. Nature. 2022 Mar 3;603(7899):138–44.

47. Utembe W, Kamng’ona AW. Inhalation exposure to chemicals, microbiota dysbiosis and adverse effects on humans. Sci Total Environ. 2024 Dec;955:176938.

48. Carmody LA, Zhao J, Schloss PD, Petrosino JF, Murray S, Young VB, et al. Changes in Cystic Fibrosis Airway Microbiota at Pulmonary Exacerbation. Ann Am Thorac Soc. 2013 Jun;10(3):179–87.

49. Fodor AA, Klem ER, Gilpin DF, Elborn JS, Boucher RC, Tunney MM, et al. The Adult Cystic Fibrosis Airway Microbiota Is Stable over Time and Infection Type, and Highly Resilient to Antibiotic Treatment of Exacerbations. Fleiszig S, editor. PLoS ONE. 2012 Sep 26;7(9):e45001.

50. Zemanick ET, Rosas-Salazar C. The Role of the Microbiome in Pediatric Respiratory Diseases. Clin Chest Med. 2024 Sep;45(3):587–97.

51. Li R, Li J, Zhou X. Lung microbiome: new insights into the pathogenesis of respiratory diseases. Signal Transduct Target Ther. 2024 Jan 17;9(1):19.

52. Lyon J. The Lung Microbiome: Key to Respiratory Ills? JAMA. 2017;317:1713–4.

53. Cole TB, Coburn J, Dao K, Roqué P, Chang YC, Kalia V, et al. Sex and genetic differences in the effects of acute diesel exhaust exposure on inflammation and oxidative stress in mouse brain. Toxicology. 2016 Dec;374:1–9.

54. da Silva Frost P. Sex Specific Mechanisms of Disease: Crosstalk Between Brain and Periphery in Inflammation [Internet]. [Riverside]: UC Riverside; 2023. Available from: https://www.proquest.com/docview/2842739982?pq-origsite=gscholar&fromopenview=true&sourcetype=Dissertations%20&%20Theses

55. Valdez JM. Systemic Inflammation Affects the CNS in an Age, Duration, and Sex Dependent Man [Internet] [Dissertation]. [Riverside]: UC Riverside; 2021. Available from: https://escholarship.org/uc/item/72d9f91x

